# Predicting Ecosystem Resilience Using Multi-Agent Reinforcement Learning

**DOI:** 10.1101/2025.06.07.658424

**Authors:** Claes Strannegård, Michał Palak, Niklas Engsner, Alice Stocco, Alexandre Antonelli, Daniele Silvestro

**Affiliations:** DEPT. OF APPLIED INFORMATION TECHNOLOGY, UNIVERSITY OF GOTHENBURG, SWEDEN; GOTHENBURG GLOBAL BIODIVERSITY CENTRE, UNIVERSITY OF GOTHENBURG, SWEDEN; DEPT. OF MOLECULAR MEDICINE AND SURGERY, KAROLINSKA INSTITUTET, SWEDEN; DEPT. OF COMPUTER SCIENCE AND ENGINEERING, CHALMERS, SWEDEN; DEPT. OF ENVIRONMENTAL SCIENCES, INFORMATICS AND STATISTICS, CA’ FOSCARI UNIVERSITY OF VENICE, ITALY; ROYAL BOTANIC GARDENS, KEW, UK; DEPT. OF BIOLOGY, UNIVERSITY OF OXFORD, UK; CAPTAIN TECHNOLOGIES LTD; DEPT. OF BIOSYSTEMS SCIENCE AND ENGINEERING, ETH ZURICH, SWITZERLAND

## Abstract

Twin models of natural ecosystems hold great promise for informing real-world decisions on sustainable land use and biodiversity conservation. However, existing simulations of animal behavior often rely on manually crafted rules, limiting their scalability and practical utility. Here, we present a flexible and scalable agent-based modeling approach that uses reinforcement learning—instead of hand-coded rules—to simulate animal behavior. We validate this approach across ten alpine ecosystems featuring wolves, chamois, and vegetation. By comparing model outputs with empirical data, we show that the simulations reproduce realistic ecological and behavioral patterns, including population dynamics, life history traits, and social interactions. We then use the model to assess ecosystem resilience under scenarios of habitat degradation, game hunting, and heat stress. Our framework paves the way for realistic simulations advancing our ability to predict ecosystem responses to disturbance and tipping points leading to biodiversity loss, in order to support conservation planning and guide the sustainable use of natural resources.

---

Ecosystems and the species they contain provide us with multiple goods and services, such as food, clean water, and material for buildings, clothes, and medicine [1]. However, more than a million species are estimated to be threatened with extinction, which means that without radical policy changes, the contribution of nature to people may soon be disrupted, with profound consequences for biodiversity and human communities throughout the world [2]. The importance and urgency of tackling the ongoing biodiversity crisis have been recognized in landmark international agreements calling for major improvements in the protection of nature worldwide [3]. Part of our ability to find solutions for this crisis is linked to the understanding of ecosystems and our capacity to predict biodiversity dynamics. Recent research has shown the potential of ecosystem models (sometimes referred to as “digital twins”) to guide the development of science-based policies that protect biodiversity while ensuring long-term human prosperity [4, 5]. Ecosystem models have also been used to study ecosystem dynamics [6] and for exploring the consequences of human activities such as recreation, farming, logging, and urban development, right from the planning stage [7], thus providing decision-makers with input for planning interventions with the least environmental impact. Additionally, ecosystem models have been employed to predict patterns of ecological interactions, such as predator-prey fluctuations and the effects of ecosystem alterations, driven by factors such as climate change and anthropogenic impact [6].

## Analytic models

Ecosystem dynamics have traditionally been described through analytic models, such as the Lotka-Volterra [8] and Arditi–Ginzburg equations [9]. These models typically describe predator-prey dynamics using systems of ordinary differential equations that are appreciated for their ability to represent population interactions over time. Their strength lies in modeling species interactions and biomass fluctuations over multiple generations, contributing to our understanding of energy and mass flow within ecosystems [10]. However, it remains difficult to model analytically the behavior of organisms in complex and more realistic scenarios, for instance, where multiple functional groups (e.g. carnivores, herbivores, and primary producers) interact, and where spatial features and environmental variables affect the dynamics of the ecosystem [11].

### Simulation models

Another approach to modeling ecosystems is based on computer simulations [12] that help solve problems for which analytic solutions are impractical or unavailable, such as ecosystems with multiple functional groups, going beyond a simple prey-predator dynamic, or that account explicitly for spatial features. Several simulation models have been developed to generate realistic representations of biological systems, including *population-based models*, which represent organisms at an aggregated level [13, 14, 4, 15], and *agent-based models*, which represent organisms individually [16, 17, 18, 19]. In agent-based models, each agent typically has a mechanism for decision-making, which governs its actions, such as moving and feeding, in a spatially explicit environment. Thanks to these characteristics, agent-based modeling has proven useful in wildlife management [20], in fishery ecology [21], and even in evolutionary contexts [15].

### Reinforcement learning

Reinforcement learning [22] is a paradigm in artificial intelligence that enables an *agent* to interact with an *environment* and learn a behavior through trial and error. The agent can interact with the environment to various degrees by making *observations* and performing *actions*. It receives feedback from the environment in the form of a *reward* signal, consisting of a number that, at each time step, quantifies to what extent the interaction between the agent and the environment has a positive or negative effect on the agent. Reinforcement learning algorithms are used to optimize models that encode how agents behave in their environment to maximize their reward. These models are called *policies*, mapping observations to actions. A common way of representing policies is to use *policy networks*: artificial neural networks that take observations as input and return actions as output [23]. Reinforcement learning has been used in domains such as robotics, autonomous driving, finance, natural language processing, and healthcare [22]. It has also been used for developing programs that can play video games like Pac-Man and Space Invaders [24], strategic games like chess and Go [25], and most recently, Minecraft [26]. In environmental sciences, reinforcement learning has been used to select optimal areas for biological conservation, where the agent is a policy maker establishing protected areas in an environment with multiple species and the reward is determined by preventing biodiversity loss [4, 27]. Additional applications of reinforcement learning within the field of ecology have been discussed [28] but have not yet led to applications for modeling real ecosystems.

*Multi-agent reinforcement learning* is a form of reinforcement learning used to develop behavioral models for multiple agents interacting within a shared environment [29]. For instance, researchers have used it to train models that play StarCraft II at a professional level [30] and to simulate predator-prey interactions in virtual maze-like environments [31, 32].

### Challenges in agent-based ecosystem modeling

In spite of the broad flexibility of simulation approaches, examples of agent-based ecosystem models remain relatively few and their potential has not been fully exploited [33]. Efficient and realistic ecosystem modeling is challenged by our limited ability to model the behavior of its organisms under a wide range of conditions. Indeed, a key component of agent-based ecosystem modeling is the encoding of animal behavior, which may include moving, feeding, and resting as a response to environmental inputs (e.g. presence or absence of predators in the vicinity) and physiological inputs (e.g. hydration and energy levels) [17]. A common approach is to hand-code animal behavior, typically by defining a set of “if-then” rules formulated based on our understanding of behavioral ecology [34]. Yet, hand-coding behavioral models is highly challenging, as the response of organisms to multiple inputs (visual, olfactory, tactile, physiological) is poorly understood in biological systems and virtually impossible to capture in an exhaustive set of rules linking all possible combinations of inputs with resulting actions [35]. One of the challenges is achieving *stable* ecosystem models, such that under beneficial circumstances, their functional groups can coexist for long periods of time in a dynamic equilibrium, without any of them dying out [36]. Another challenge in hand-coding behavioral models is obtaining real-world behavioral data, specifying in detail how individual animals act in different situations that they may encounter in their natural environments. This is also a limiting factor for developing behavioral models based on supervised learning [37]. Yet another challenge related to agent-based ecosystem modeling is obtaining population data for all functional groups, which might be used to define starting points of simulations and to validate the models. We propose to tackle one of the main challenges of agent-based ecosystem modeling by using reinforcement learning rather than hand-coding for constructing models of animal behavior. This is a special case of a more general idea—replacing rule-based models by machine learning models—which has been highly successful in other scientific areas, such as predicting protein structure [38] and, most recently, weather forecasting [39].

## Results

We developed a new general strategy for constructing agent-based ecosystem models powered by multi-agent reinforcement learning and used it to create models of real ecosystems and explore to what extent these models were able to reproduce realistic ecosystem dynamics and animal behavior. Our ecosystem models were based on ten alpine areas, located in the Rhaetian Alps around the Stelvio National Park, in Italy (Fig. 1(A)). An overview of the flora and fauna of the Rhaetian Alps can be found in [40, 41, 42]. Our ecosystem models included: (1) primary producers represented by three types of vegetation, each with its own growth rate; (2) a large herbivore, hereafter referred to as the chamois (*Rupicapra rupicapra*), one of the mammalian herbivores in the region; and (3) an apex predator, hereafter referred to as the wolf (*Canis lupus*), one of the top mammalian predators in the area [42]. The primary producers were modeled collectively, while the chamois and wolves were modeled individually as agents. In the Alpine area, chamois and wolves have been able to coexist for hundreds of years. After a period when wolves were hunted to extinction, migrations and human reintroduction repopulated the area [43, 42]. We used geographical information to construct *spatial models* of the ten areas, discretized into 100 *×* 100 cells and populated with vegetation and agents representing individual chamois and wolves (Fig. 1(B)). After defining a *dynamic model*, which specifies the rules for reproduction, death, and the consequences of the agents’ actions, we used reinforcement learning to train a *behavioral model*, which computes the action of any agent at any location in any spatial model. The behavioral model was trained by using thousands of randomly populated “synthetic” spatial models, generated using Perlin noise [44] (Fig. 1(C)–(D)). The purpose of using multiple spatial models was to produce versatile behavioral models of chamois and wolves that can cope with a range of different environments. We then used our trained behavioral models to conduct experiments, allowing us to highlight different properties of our ecological models across several areas.

**FIGURE 1.**
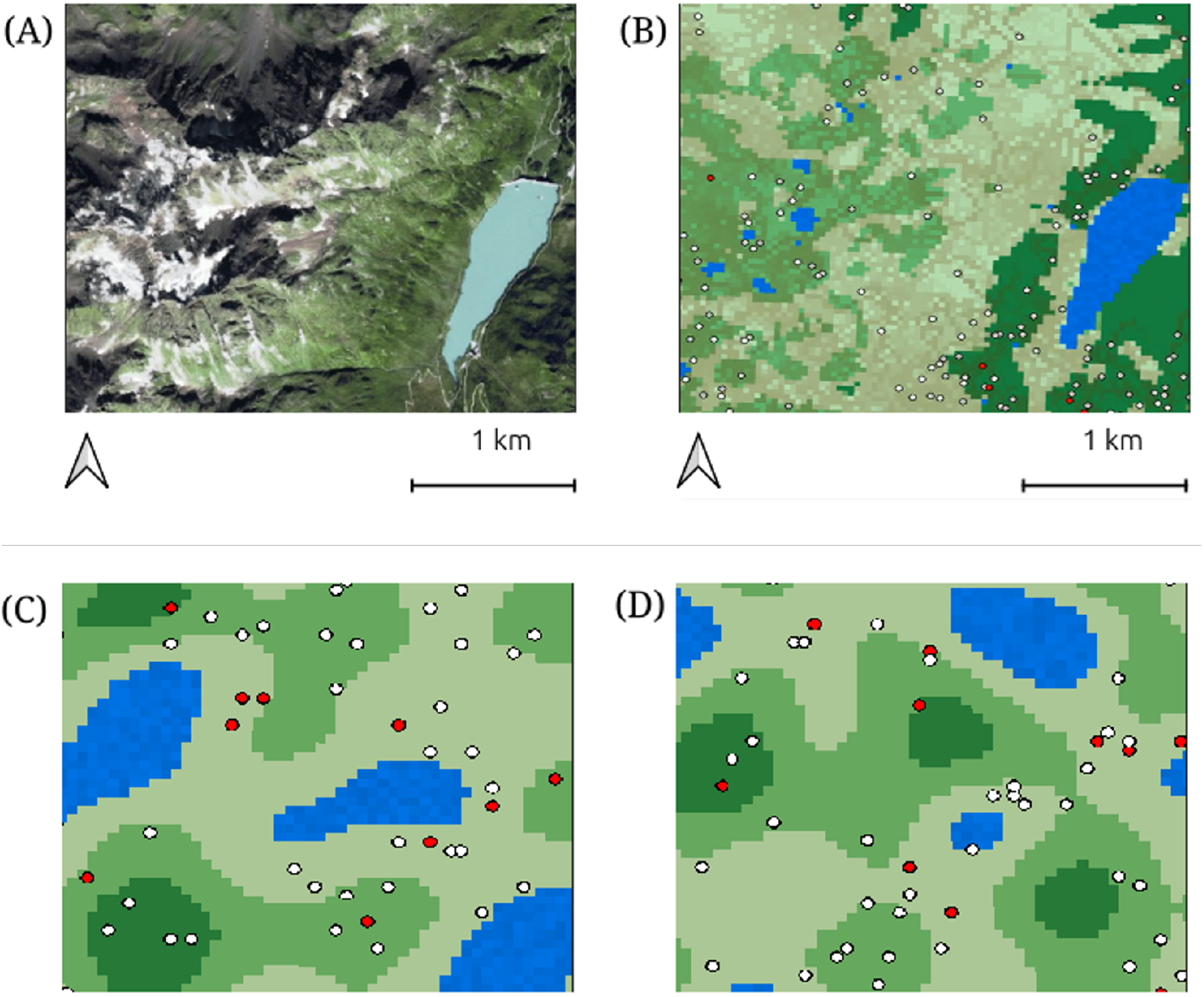
(A): Aerial photograph from 2020 of an area in the Rhaetian Alps. (B): A spatial model of the same area with 100 *×* 100 cells. Blue represents water, while the three shades of green represent areas with three types of vegetation, with darker green representing vegetation that grows relatively fast. The white and gray dots represent chamois of different age, while the red dots represent wolves. Brown indicates vegetation cells that have recently been grazed by chamois. (C) and (D): Two of the many thousand synthetic spatial models with 50 *×* 50 cells that were used for training the behavioral model.

### Coexistence experiments

The purpose of these experiments was to see to what extent the chamois and wolf agents (hereafter, the chamois and wolves for simplicity) were able to coexist in spatial models of the ten areas of the Rhaetian Alps (Fig. 2). Ten simulations were run on each of the ten spatial models, starting from randomly spawned populations. Thus, there was a total of 100 simulations. Each simulation was run for up to 4,000 steps, corresponding to eight times the maximum lifetime of a wolf (which was set to 500 steps), but it was stopped as soon as an extinction event occurred (either the chamois or the wolves went extinct). In 95 of these 100 simulations, neither the chamois nor the wolves died out. In the remaining five simulations, the wolves died out (Supplementary Fig. 3). These simulations showed that the agents with behavior optimized through reinforcement learning had developed the ability to coexist in spatial models of the ten natural environments.

**FIGURE 2.**
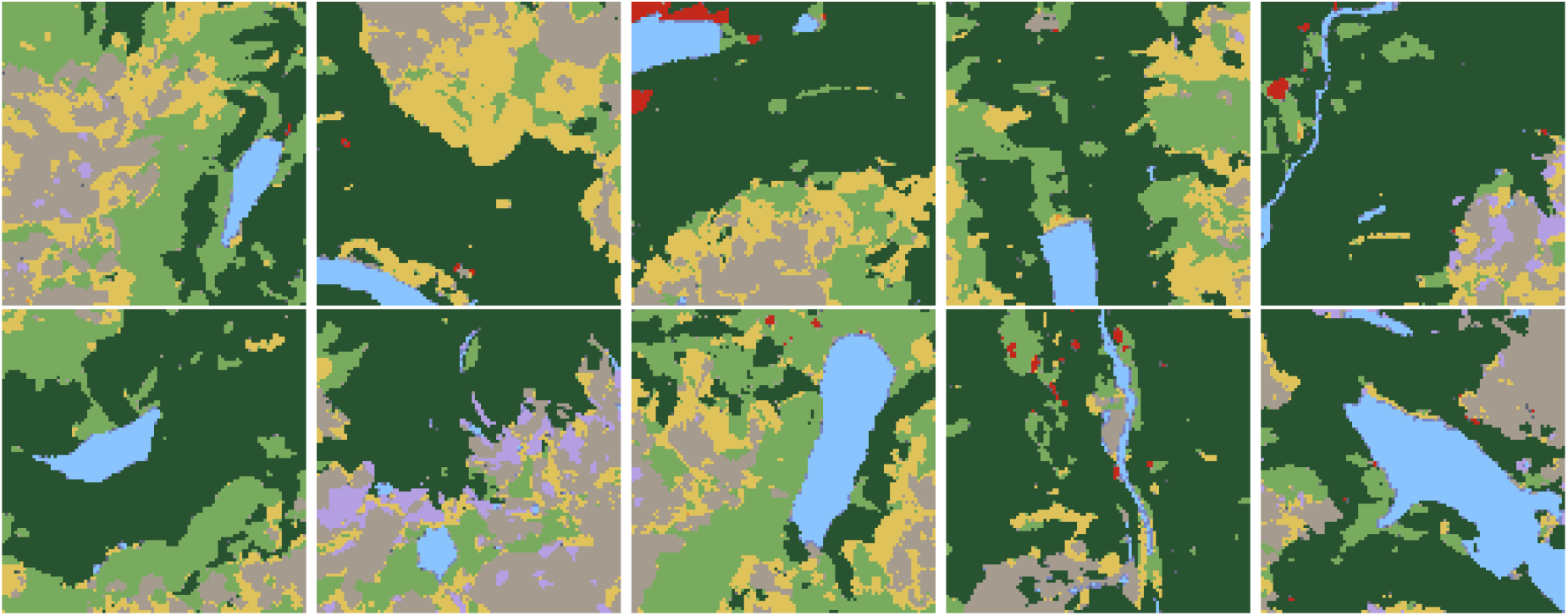
Maps of ten areas in the Rhaetian Alps based on multi-band Sentinel-2 images with 100 *×* 100 pixels. Light blue represents water, gray is used for rocks and bare soil, lilac stands for snow, yellow for scarcely vegetated areas, light green for pastures and grasslands, dark green for shrubs and tree-vegetated areas, while red is used for buildings and artificial settlements.

**FIGURE 3.**
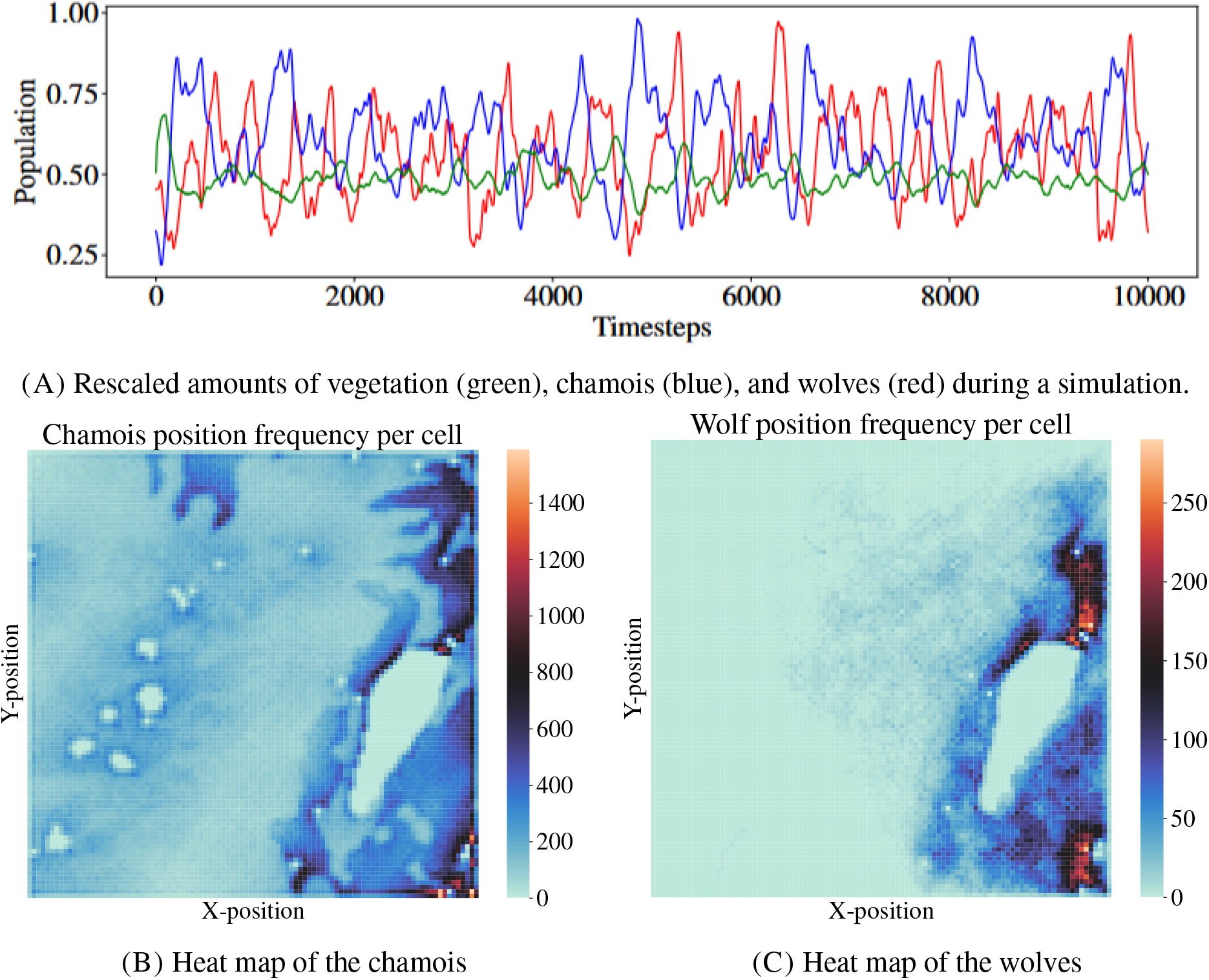
Population dynamics from the behavioral experiment. (A): The graph shows the amount of wolves, chamois, and vegetation during the simulation run. For each of these functional groups, the amount was normalized to the interval [0, 1]. The max level 1.0 represents 35 wolves, 267 chamois, and fully grown vegetation across the map. (B) and (C): Heat maps for chamois and wolves, reflecting their spatial distribution during the simulation.

### Behavioral experiment

We then ran an experiment to evaluate the ability of the framework to reproduce and maintain long-term realistic population dynamics and animal behavior, in the absence of external disruptive factors. Using one of the ten spatial models (Fig. 1(B)), we ran a simulation for 10,000 steps, which corresponds to 20 times the maximum lifetime of a wolf. The population dynamics observed in this experiment show that the chamois and wolves were indeed able to coexist for multiple generations in the same area. The number of agents alive at the same time during the simulation ranged between 64 and 307 for the chamois and between 7 and 33 for wolves. These numbers reflect the expected imbalance between the abundances of prey and predators [45, 46]. Moreover, Lotka-Volterra dynamics arose as an emergent property of the model. Our simulation revealed a clear and consistent pattern where a chamois population surge is followed by a wolf surge, which leads to a chamois decline, followed by a wolf decline, before the cycle starts over again (Fig. 3(A)). This is remarkable because such dynamics had not been manually encoded in the behavior of the agents, but emerged from the simulation. The location of the chamois and wolves agents was monitored during the simulation so that heat maps of their spatial distribution could be constructed. The heat maps indicate that the chamois were relatively spread throughout the area (Fig. 3(B)), while the wolves were found mainly around the big lake (Fig. 3(C)). The statistics collected show that many chamois died at a young age and only a few reached the maximum age (Fig. 4(A)). Young wolves had high death rates, but those that survived past early life tended to live relatively long lives (Fig. 4(B)). A comparison of these statistics (Fig. 4) against available data about natural populations of chamois and wolves in the literature showed that our simulations led to realistic patterns [43]. The animal agents in the simulation can seem to behave in a similar way to their natural counterparts (Supplementary Video 1). Chamois tend to move between vegetation and water sources; wolves tend to hunt chamois, and chamois tend to run away from wolves; both chamois and wolves tend to move along relatively straight trajectories, while avoiding obstacles such as lakes; and both chamois and wolves have a tendency to form groups.

**FIGURE 4.**
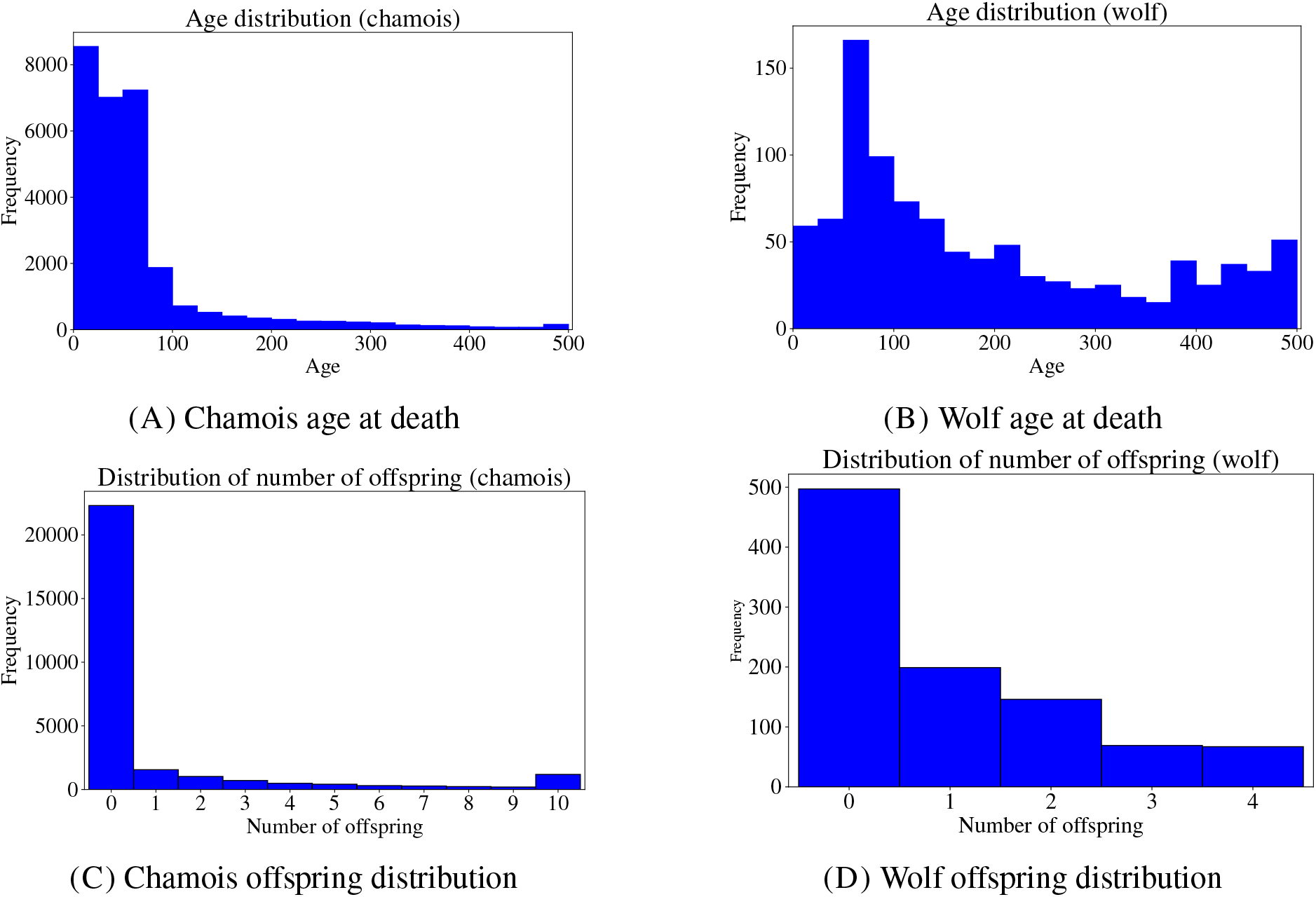
Life history statistics from the behavioral experiment. Longevity distribution of (A) chamois and (B) wolves. Offspring distribution of the chamois (C) and wolves (D). The rightmost bar in plot (C) represents ten off-spring or more.

### Resilience experiments

The purpose of these experiments was to evaluate the resilience of the ten ecosystems (Fig. 1) to various degrees of environmental pressure in the form of habitat degradation, game hunting, and heat stress. In particular, we wanted to see if we could locate tipping points, where the pressure would likely cause a decrease in biodiversity.

#### Habitat degradation

Habitat degradation—potentially caused by agriculture, forestry, mining, infrastructure projects, or urban development—was simulated by replacing areas of fast-growing natural vegetation with slow-growing vegetation. This effectively reflects a transition from a naturally productive pasture to one that take substantially longer time to recover its full biomass after grazing. Thus, habitat degradation led to more limited resources for the chamois. We explored the effect of different levels of habitat degradation, ranging from 0% to 100% of the area covered with vegetation. Each level of habitat degradation was tested on ten different spatial models and each model was simulated ten times using randomized initial animal populations. Thus, we ran 1,100 simulations in total. The simulations indicate a tipping point at *∼* 40% of habitat degradation (Fig. 5(A)).

**FIGURE 5.**
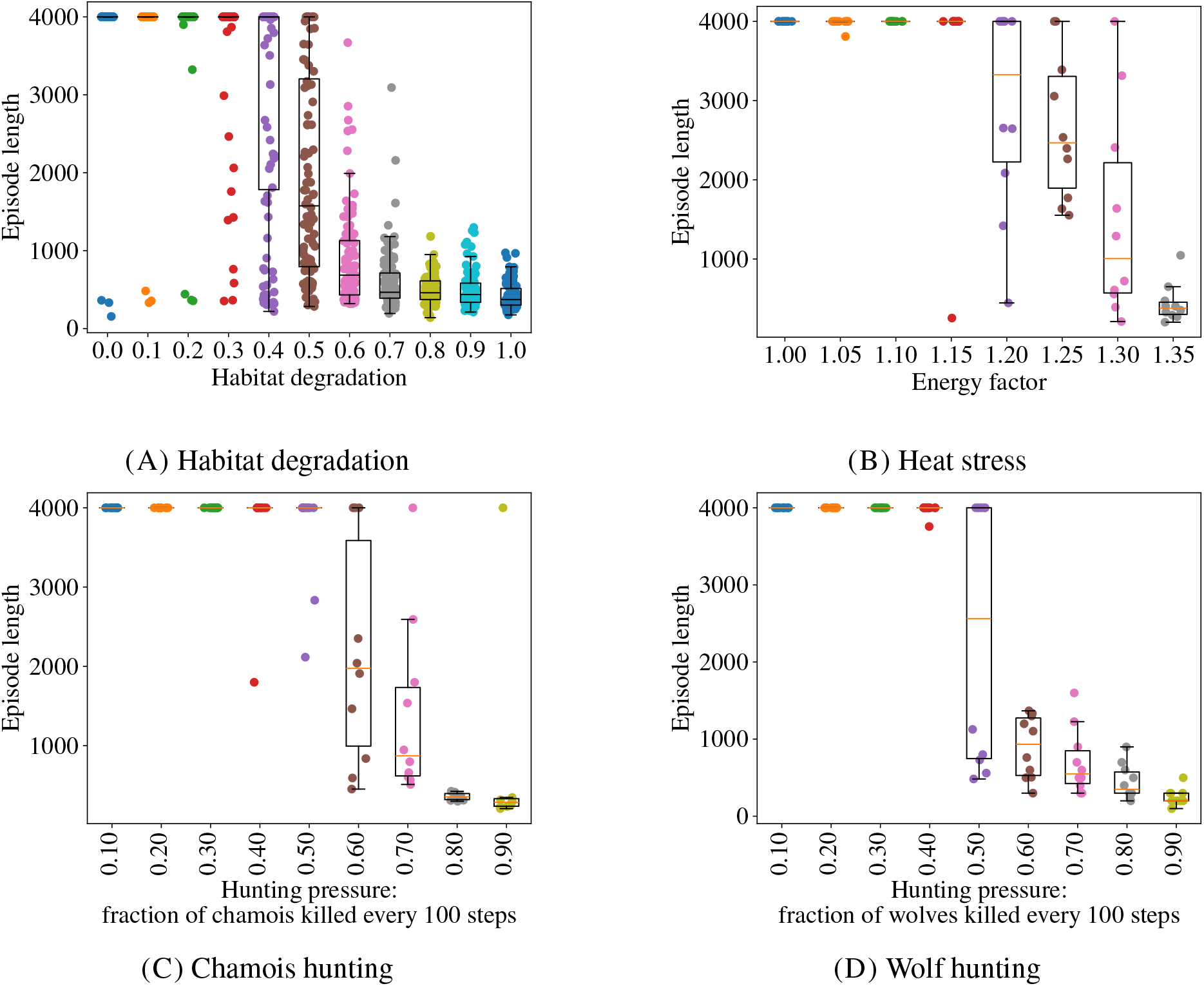
Results from our resilience experiments. Each colored dot represents a simulation and its y-coordinate indicates how long the simulation lasted. Each simulation (episode) was run for up to 4,000 steps, but it was stopped as soon as an extinction event occurred. Our experiments studied the episode length for different levels of (A) habitat degradation; (B) metabolic energy consumption; (C) chamois hunting; and (D) wolf hunting.

#### Game hunting

To model game hunting, we randomly removed animals, at nine levels of hunting pressure (ranging from 0% to 90% killed every 100 time steps). Wolf hunting and chamois hunting were studied separately. We ran ten simulations at each level of hunting, thus generating 90 data points for the chamois and 90 for the wolves. The simulations indicate that there are tipping points for hunting at, respectively, *∼* 50% for chamois and *∼* 40% for wolves (Fig. 5(C) and 5(D)), above which an extinction event occurred. In all simulations where extinction events occurred, the wolves died out first, even when hunting targeted the chamois only. This reveals a higher sensitivity of the predator species in our system to increased mortality.

#### Heat stress

We explored the effect of prolonged heat stress, for instance resulting from extreme weather patterns linked to climate change. Increased temperature puts pressure on many animal species by causing their energy consumption (metabolic rate) to increase. We studied the effect of heat stress at eight different levels of increased energy consumption (in the range 0–35%) for both chamois and wolves. Each level of stress was simulated ten times, starting from the same initial environment in all cases. Thus, a total of 80 simulations were performed. In the simulations, there was virtually no effect of increased temperature up to a tipping point at (*∼* 20%), after which the extinction risk increased sharply (Fig. 5(B)). In all simulations with extinction events, the wolves died out first.

## Discussion

We have constructed a highly flexible strategy for ecosystem modeling that captures animal behavior and population dynamics without the need for hand-coding behavioral rules. This constitutes a first step towards robust exploration of the impact of individual interventions – such as habitat degradation, selective hunting or ecological restoration – on ecosystem health, integrity and biodiversity.

### Realism

Real ecosystems may contain thousands of species, millions of individuals, and countless known and unknown mechanisms that govern animal behavior and ecosystem dynamics. How ecosystems function and respond to changes depends strongly on phenomena that are not fully understood and commonly modeled as stochastic processes, for example reproduction, death, migration, genetic mutations, and climate and landscape changes [47]. Because of this complexity, ecosystem models must inevitably rely on simplifying assumptions, for example when defining food webs. The simplifying assumptions must be chosen with the intended use of the ecosystem model in mind, whether it is to explore ecological patterns, analyze different management interventions, or predict resilience and ecological tipping points. The simplifying assumptions that were used in our models include a food web focused on a few representative species within trophic groups, imposed area boundaries that do not account for migration, and a reproduction model that does not fully capture biological behaviors. Yet, our results showed that the models were able to reproduce realistic ecological and behavioral patterns including population dynamics, life history statistics, foraging, and social behavior.

### Scope

The behavioral and dynamic models used in our experiments can be trained to model different ecosystems in any geographical area. This gives the model a considerable flexibility in its biogeographic scope and the biological species it can include. Our modeling framework can be extended with the addition of functional groups (for instance scavengers or omnivores) that can make the food web more complex and realistic. Additional land cover classes (e.g., more vegetation types) can also be included in the ecosystem model. After defining the properties of the different species and vegetation types the ecosystem models can be trained through reinforcement learning following the procedure utilized in our experiments. There are limits to the number of agents that can be included in agent-based models, depending on the time available for training and inference, and the available computational resources. We used hundreds of agents in our experiments, while millions of agents have been used in other studies based on reinforcement learning [31], leaving substantial room for scaling up the scope of our ecosystem models.

### Perception

Our animal agents receive input signals that can be divided into internal signals (interoception) and external signals (exteroception). The internal signals inform the agents about their status, in this case with regard to energy and hydration, while the external signals inform the agents about the quantity and location of food sources, water sources, competitors, predators, and obstacles in their surrounding environment. In reality, such information can originate from a combination of multiple senses, possibly together with memories of earlier observations. The external signals can be divided into short-range and long-range signals. The short-range signals provide relatively precise quantitative information about each cell in the closest neighborhood surrounding the agent. The long-range signals, which we call “smells” for simplicity, provide the agents with less precise, aggregated information about cells outside their closest neighborhood. By perceiving and comparing the smell intensity in the nine cells surrounding it, the agent obtained approximate information about the amounts and direction of the sources of the smell. Thus, smell might help agents locate food, water bodies, competitors, and predators.

### Emergent behavior

During training, the animal agents were directly rewarded for eating and drinking. Indirectly, they were also rewarded for surviving, since surviving enables them to eat and drink more in the future and thus receive a greater total reward. Thus, the agents were trained to eat, drink, and survive in a broad range of environments, just like real animals are able to do. Our results from the coexistence experiment indicate that the trained agents are indeed able to survive in a range of spatial models based on real geographical data. This is not an obvious outcome, since the agents were never trained on empirical landscape models, but only on synthetic environments. For the chamois, surviving means being able to obtain food and water, while efficiently navigating in the presence of obstacles and actively escaping predation from wolves. For the wolves, it means being able to locate, chase and hunt chamois. These behaviors, which the agents share with their natural counterparts, were also observed in our behavioral experiment (Supporting Video 1). Just like real animals, our animal models were able to coexist for multiple generations in a range of different environments. This means that they had developed an ability to survive and reproduce by tackling the stream of challenges that their environment poses to their cognitive abilities. The behavioral strategies that emerged during training, along with the biologically realistic population dynamics, show that this approach based on reinforcement learning is promising to help overcome the challenges of modeling complex social-ecological systems.

### Conclusions

Here we have shown how to tackle a main difficulty in agent-based ecosystem modeling by using machine learning rather than hand-coding for modeling animal behavior. We used this strategy to construct ecosystem models of ten different areas in the Rhaetian Alps. In these ecosystem models: (i) the chamois and wolves were able to coexist for long periods of time without any of them dying out; (ii) several characteristic patterns of animal behavior and population dynamics emerged; and (iii) ecological tipping points could be identified when the models were exposed to increasing levels of habitat degradation, game hunting, and heat stress.

Our results indicate that it is possible to build agent-based ecosystem models that leverage machine learning and that the resulting ecosystem models are useful for analyzing and comparing different management policies from the perspective of ecology and sustainability. Next steps in this research might include building and validating more detailed models of ecosystems where high-resolution time-series data about animal behavior and population dynamics exist, and introducing additional modeling dimensions such as sexual and seasonal reproduction, altitude, and temperature. As ecosystem twins increase in popularity, we hope that behavioral models powered by machine learning will pave the way for more realistic, scalable, and useful ecosystem models for policymakers and scientists alike.

## Methods

### Modeling strategy

We used a general strategy for constructing agent-based ecosystem models that combined three key ideas: (i) to model animals as reinforcement learning agents that make decisions based on their perception of the environment and their internal homeostatic signals; (ii) to reward agents for living long lives and thus, indirectly, for eating and drinking, avoiding predators and other lethal dangers, and navigating efficiently in the terrain; and (iii) to train the agents in many different environments so that they become flexible enough to survive in models of many different geographical areas with varying ecological conditions. Before describing in detail how our ecosystem models were developed, we outline our general strategy for ecosystem modeling.

#### Conceptual model

Given a set of ecosystems and ecological phenomena that one wishes to model, we first defined the building blocks, or the ontology, of our ecosystem models. A *conceptual model* consists of (i) a set of *functional groups*, which is divided into decision-making and non-decision-making functional groups, where decision-making functional groups (e.g., wolves and chamois) are modeled individually as agents, and non-decision-making functional groups (e.g., different kinds of vegetation) are modeled collectively; (ii) a set of *agent properties*, for example age, weight, max speed, max age, energy level, hydration level, position, observation space, and action space; (iii) a set of *cell properties*, for example, biological properties such as the abundance of each functional group, physical properties such as altitude and temperature, and landscape properties such as land-cover class (with values like rock, field, sand, water). The cell properties might also include smell intensity of different functional groups and (organisms associated with) land-cover classes.

#### Spatial model

We use spatial models to represent ecosystems at a given moment in time. A *spatial model* consists of a grid of *cells*, where each cell has its own cell properties, and a set of *agents*, with its own agent properties (Fig. 1(B)).

#### Behavioral model

A *behavioral model* consists of the following components, for each decision-making functional group: (i) an *observation space*; (ii) an *action space*; and (iii) a *policy network* computing a function from the observation space to the action space. We use one policy network per decision-making functional group for computing the actions of each agent, based on the properties of the cells surrounding it and its own internal state (Fig. 6(A)).

**FIGURE 6.**
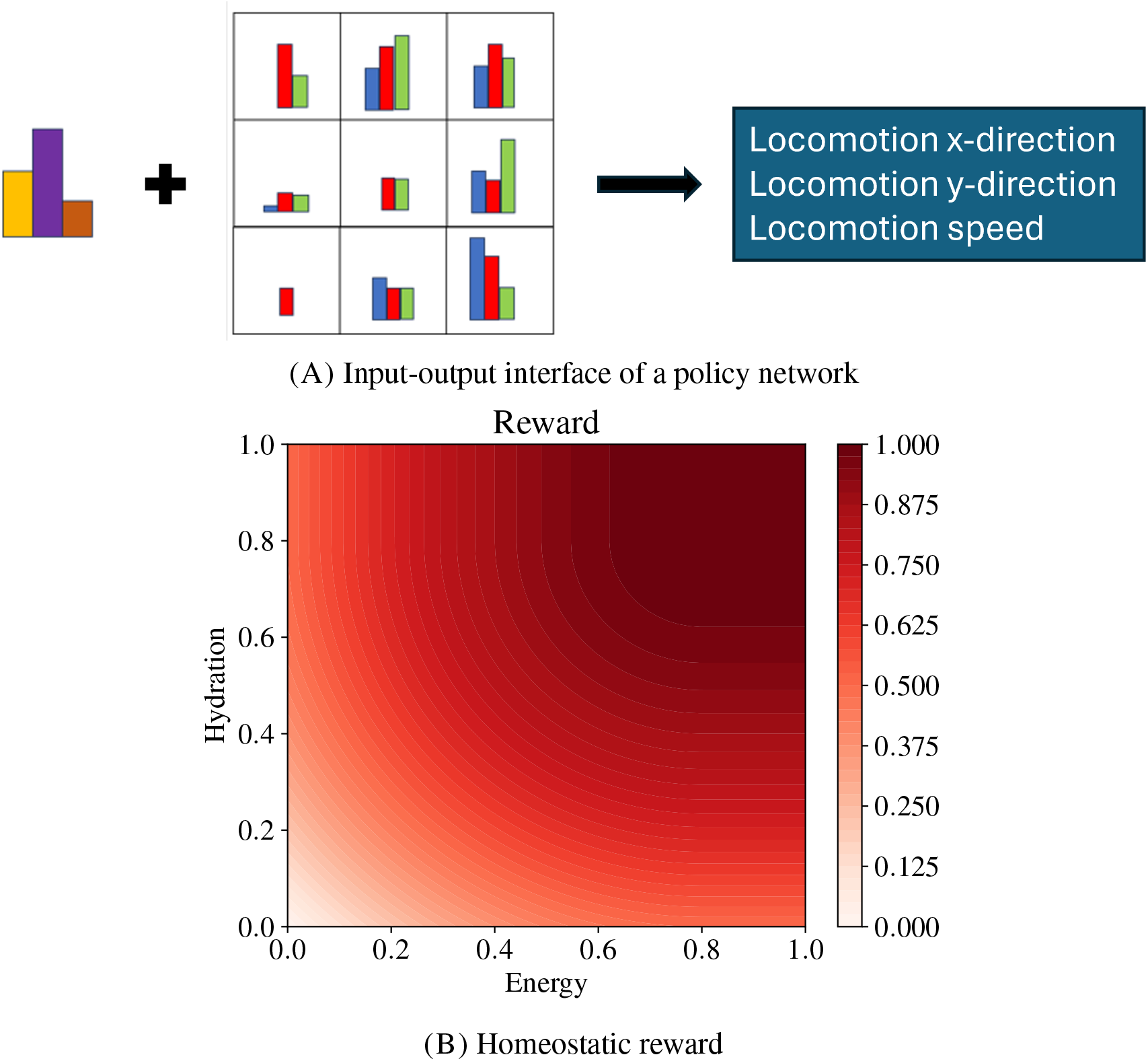
(A) Interface of a shared policy network between chamois and wolf. The agent observes its own internal state, including its own functional group (left) and the cell properties of the 3 *×* 3 cells surrounding it (middle). Thus, the agent can perceive its own state, along with the presence of resources, obstacles, predators, and competitors in its near environment. Given this input, the agent uses a policy network to compute its movement, given by a direction and a speed (right). (B) The homeostatic reward used for training policy networks, quantifying the “well-being” of the agent in terms of its energy and hydration levels. An agent with both energy and hydration levels above the saturation threshold of 0.8 will not get additional reward from increasing its energy or hydration levels further.

#### Dynamic model

A *dynamic model* is a mechanism that takes as input a spatial model and the decisions (actions) of all agents of that spatial model; and outputs an updated spatial model. Dynamic models are typically defined as a set of update rules specifying how the primary producers grow, the consequences of the agents’ locomotion and feeding actions, and how the physical properties of the cells develop over time.

#### Training process

Given a conceptual model *C* and a dynamic model *D*, we use reinforcement learning to construct a behavioral model *B*. More precisely, we fix an architecture for the policy network of each decision-making functional group and train their parameters (weights and biases) with reinforcement learning. Thus, a reward signal is used, which provides an assessment of the impact of the agent’s actions. For instance, actions leading to improved energy levels are typically assigned a positive reward, while actions leading to the death of an agent is assigned a negative reward. The goal of the training process is to optimize the parameters of the policy network toward maximizing the cumulative reward in the long run. The policy networks of all decision-making functional groups can be constructed simultaneously as follows: (1) Define a class of spatial models *ℒ* that are suitable for training; (2) Initialize the behavioral model *B* by using randomization; (3) Repeat the following until *B* is good enough: For every spatial model *S ∈ ℒ*, use *B* and *D* to run a simulation starting from *S*, for a fixed number of steps, then update *B* using reinforcement learning. The reason for training the behavioral model *B* on a class of spatial models *ℒ* rather than a single spatial model is to achieve generalization properties [48]. When defining *ℒ*, one may consider using synthetic data, which have been used successfully in supervised learning as well as reinforcement learning, e.g., when data are scarce [49].

#### Generic model

Finally, we put the pieces together and define a *generic model*, comprising three elements: a (1) conceptual model, (2) dynamic model, and (3) behavioral model. With a generic model, one may start with any spatial model based on the same conceptual model and then apply the behavioral and dynamic models repeatedly, to generate a sequence of spatial models that simulate how ecosystems develop over time.

### Alpine model

Next we show how we used the general strategy described above to construct a generic model for alpine ecosystems.

#### Conceptual model

Our conceptual model included two decision-making functional groups (chamois and wolf) and three non-decision-making functional groups (three types of vegetation). We only used two land-cover types: land and water. The cell properties included biomass and smell of the functional groups, and smell of water. For both chamois and wolf agents, the observation space consisted of two internal variables (energy and hydration) and the cell properties of a 3 *×* 3 cell neighborhood, while the action space consisted of triplets of numbers encoding direction and speed.

#### Behavioral model

For simplicity we let all agents belonging to the same functional group share the same behavioral model. Thus, the decisions of all chamois are taken by the same policy network and similarly for the wolves. Note that different agents will usually perceive the environment differently, based on their current location, so the actions computed by the policy network will often differ among them at any given time. Moreover, to simplify the behavioral model further, we used a single policy network for both chamois and wolves. To that end, we added an input to the policy network for encoding the functional group (chamois or wolf). The input to the policy network also included the agent’s other properties and the properties of the 3 *×* 3 cells surrounding it, all in all 144 inputs. The output of the policy network was a triplet of real numbers (Δ*x*, Δ*y, v*), specifying movement in the direction (Δ*x*, Δ*y*) at speed *v*. The speed *v* is a number in [0, 1] that specifies a fraction of the agent’s maximum speed. The network had a fully connected architecture with two hidden layers with 64 nodes each and with tanh as the activation function (Fig. 6(A)).

#### Dynamic model

We now describe how the set of agents, the agent properties, and the cell properties were updated at each time step.

Primary production. When a chamois was on a cell, the biomass of the vegetation of that cell was set to 0, modeling grazing. Then the vegetation grew back gradually, reaching a predefined maximum biomass after 20, 100, or 200 steps, depending on the vegetation type.

Metabolism. A chamois in a cell containing vegetation would graze automatically. Thus, the energy level of the chamois was increased by up to 3% of its maximum energy level, while the vegetation disappeared and started to regrow at a speed that depended on its type. Similarly, a wolf in the same cell as a chamois would kill the chamois automatically. Thus, the energy level of the wolf was increased by up to +18% of its maximum energy level. Moreover, both chamois and wolves drank automatically every time they were in a cell adjacent to water, thus reaching a hydration level of 100%.

Movement. The agents could use their actions to move, rest, feed, and drink. In fact, resting corresponds to moving at speed 0, while eating and drinking happened automatically, as mentioned. Although the space was divided into a discrete grid of cells, we kept track of the exact position of each agent in continuous space, by including the coordinates (*x, y*) *∈* R_2_ of its position among the agent properties. The agents could move in any direction and at different speeds, as determined by the output of the policy network. The maximum speed varied with the age of the agent and was set to increase linearly until an age of 100 time units, after which it plateaued until 250 units, whereupon it declined linearly until 500 units (i.e., the maximum age). We set the maximum speed of the wolf to slightly exceed that of the chamois. The rule for updating the position of an agent was as follows: If the position of an agent at time *t* is (*x*_*t*_, *y*_*t*_) and the output of the policy network of the agent is (Δ*x*, Δ*y, v*), then its position (*x*_*t*+1_, *y*_*t*+1_) at time *t* + 1 will be (*x*_*t*_ + Δ*x · v, y*_*t*_ + Δ*y · v*).

Reproduction. Both chamois and wolves reproduced with a probability that was set to 0.1, given that their energy level was above a threshold, here set to 95% and 50% of their maximal energy level for wolves and chamois, respectively.

Death. Individual chamois and wolves would die from starvation if they ran out of energy, from thirst if they ran out of hydration, from drowning if they entered a water cell, and from old age if they reached their predefined maximum age, which was set to 500 steps. In addition, the chamois would die from predation if they were caught by a wolf.

Smells. Once the other cell properties had been updated, we computed the smells. We used a basic concept of smell. For example, to compute the smell of wolf or water in a certain cell, we simply took the Euclidean distance to the closest wolf or closest water cell.

#### Training process

Reinforcement learning was used to train the policy networks of the chamois and the wolf. We used a *homeostatic reward* signal for each agent, which measures its “well-being”, defined as a quadratic function of its energy and hydration levels (Fig. 6(B)). Thus, the policy network was optimized to keep the energy and hydration levels of each agent above a threshold that was set to 80% of the maximum energy and hydration level, respectively. The policy network was trained with reinforcement learning using the PPO algorithm from Stable-Baselines3 [50]. For training, we created thousands of synthetic spatial models by using Perlin noise [44], which is widely used in the movie and gaming industries for generating artificial landscapes. More precisely, the spatial models were created by first defining a landscape with 50 *×* 50 cells using Perlin noise and then randomly spawning 40 chamois and 10 wolves on that grid (Fig. 1(E)–(F)). At the start of each training episode, a new spatial model (Perlin world) was constructed using randomization. By training on many Perlin worlds, the agents were encouraged to learn general behavioral rules that could be useful in many spatial models, instead of learning the specifics of a single spatial model. Again for the sake of efficiency, the number of agents was kept constant during training by automatically adding a new individual of the same functional group, when a death occurred. We used intermediate checkpoints of the model obtained during training to monitor their ability to maintain stability, where none of the functional groups goes extinct. During validation, episodes were run for 4,000 time steps, but they were aborted as soon as one of the functional groups (specifically the chamois or wolf) went extinct. More information about the training process, including the choice of algorithm, reward signal, and hyperparameters can be found in the Supplementary Information.

### Spatial models

We used land-cover maps obtained by classifying multiband satellite images, retrieved in 2023 by Copernicus Sentinel-2 fleet, of ten non-overlapping 4 *×* 4 km areas in the Rhaetian Alps. All ten contain water bodies, as the basis for our spatial models (Fig. 2). To create spatial models from these maps, we populated them by the three types of vegetation, which were distributed following the land cover types and initially set to 50% of their maximum biomass in each cell. Moreover, 100 chamois and 25 wolves were randomly spawned on the land cells of the map. The agents were spawned randomly since—as far as we are aware—no high resolution population data exist for the ten areas considered.

## Data availability

All data supporting this research are available in the main text and the Supplementary Information. Original data can be obtained from the corresponding authors upon request.

## Code availability

The code of the ecosystem models is implemented in Python v.3 and is available in the repository https://gitlab.com/ecotwin/abm_paper_2024.git.

## Acknowledgments

CS acknowledges funding from the Sten A Olsson Foundation for Research and Culture, Stiftelsen Sävs-taholm, and Erik och Lily Philipsons Stiftelse. DS received funding from ETH Zurich. AA and DS acknowledge financial support from the Swedish Research Council (2019-05191; 2024-04303) and the Swedish Foundation for Strategic Environmental Research MISTRA within the framework of the research programme BIOPATH (F 2022/1448). AA also acknowledges funding from RBG Kew Development. The computations were enabled by resources provided by the National Academic Infrastructure for Supercomputing in Sweden (NAISS), partially funded by the Swedish Research Council through grant agreement no. 2022-06725. We thank Rhian Smith for feedback on this manuscript.

## Notes

### Competing Interest Statement

CS and DS are shareholders in Ecotwin Sweden AB. AA and DS are shareholders in Captain Technologies Ltd.

