## Supplementary material for "Predicting Ecosystem Resilience Using Multi-Agent Reinforcement Learning": Supplenentary Information

### SUPPLEMENTARY METHODS

**Training process.** The training process used the reinforcement algorithm PPO from the repository Stable-Baselines3 [50]. It spanned 4 million time steps and took about 8 hours on a standard laptop computer. Each training episode lasted 2,000 time steps. The models were tested repeatedly during training by measuring the average episode length over several test runs until some functional group died out. Supplementary Table 1 shows the hyperparameter settings that were used when training the policy network.

| **Hyperparameter** | **Value** |
|---|---|
| `learning_rate` | 0.0003 |
| `n_steps` | 512 |
| `batch_size` | 64 |
| `n_epochs` | 10 |
| `gamma` | 0.99 |
| `gae_lambda` | 0.95 |
| `clip_range` | 0.2 |

SUPPLEMENTARY TABLE 1. Hyperparameters used when training the policy network.

**Reinforcement learning algorithm.** To determine which reinforcement learning algorithm to use for training our policy network, we conducted a simple study. We studied three policy networks with two hidden layers each, trained, respectively, with the algorithms PPO, TRPO, and TQC, from Stable-Baselines3 [50]. We also included a fourth policy network with no hidden layers, which was trained with PPO, referred to as PPO2 [1]. After training, the four policy networks were tested on ten spatial models generated with Perlin noise. For each policy network, a 4000 step simulation was run on each spatial model. Among a total of 40 simulations, all but five lasted the maximum 4000 steps (Supplementary Fig. 1). We also studied the four policy

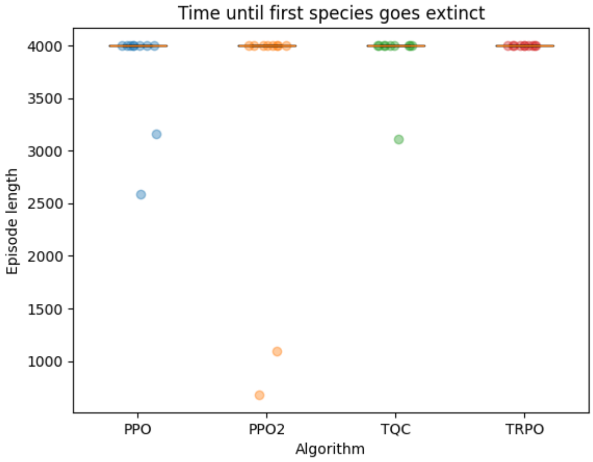


SUPPLEMENTARY FIGURE 1. Left: Episode length for different reinforcement learning algorithms on ten previously unseen Perlin worlds.

networks by running a relatively long simulation, where chamois and wolf agents with different (heritable) policy networks interacted with each other. In the simulation, the PPO2 agents died out first. (Supplementary Video 2). Against the background of these studies, we decided to go with the standard PPO algorithm.

**Reward signal.** To determine which reward signal to use for training our policy networks, we conducted a mini-study, comparing the *homeostatic* reward (Fig. 6(B)) and the *survival* reward, which gives a reward of +1 for each step, as long as no functional group has died out, cf. [2] (Supplementary Video 3). Essentially, the homeostatic reward encourages eating and drinking, while the survival reward encourages long-term coexistence of the functional groups. Our mini-study showed that the ability of the functional groups to coexist in Perlin worlds improved over time, for both reward signals (Supplementary Fig. 2). The performance seems to culminate after about 2.8 million steps for both reward signals. A possible reason for this similarity could be that the homeostatic reward implicitly encourages long lives, since agents that avoid predators

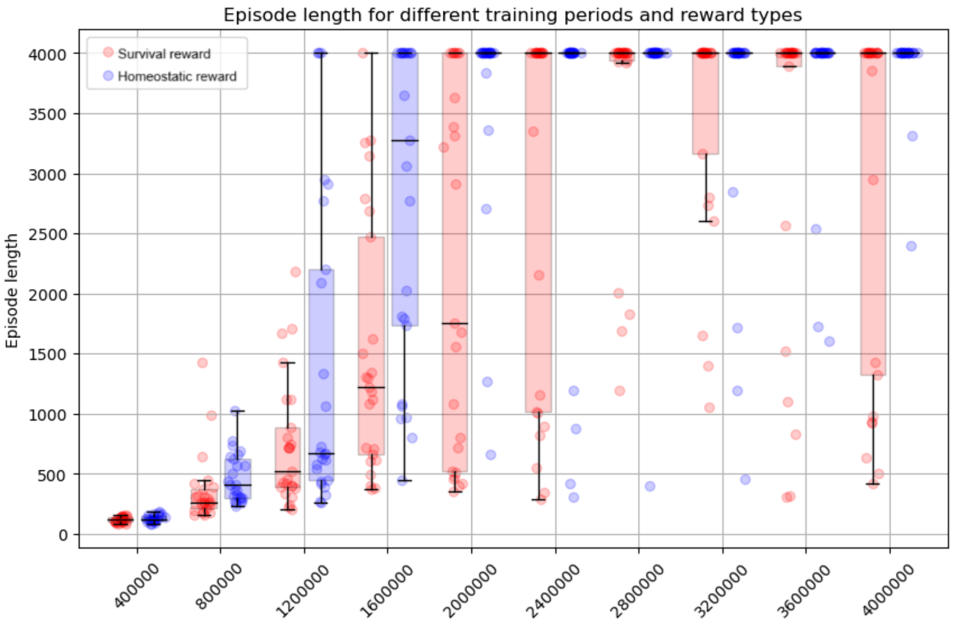


SUPPLEMENTARY FIGURE 2. Episode length during training with the homeostatic reward (blue) and survival reward (red). The x-axis shows the total training time in steps. Each dot corresponds to a test run, which lasted a maximum 4,000 steps and was aborted as soon as some functional group went extinct. The median episode length, marked with a horizontal bar, eventually reached the maximum 4,000 time steps with both reward signals.

and other lethal dangers will generally be able to eat and drink more during their lives and thus receive more homeostatic reward. Conversely, survival reward implicitly encourages behavior such as eating, drinking, predator avoidance, and navigation efficiency, since those behaviors tend to be associated with survival. We decided to use the homeostatic reward, since it led to somewhat faster and more robust learning. On the other hand, the survival reward has a great advantage in terms of generality, since it requires no reward engineering (like defining a particular weighted sum of energy and hydration) and can be used for all functional groups. For example, an anteater model might spontaneously learn to eat ants, whereas a hummingbird model might learn to drink nectar—both without any reward engineering whatsoever.

### SUPPLEMENTARY NOTES

Supplementary Fig. 3 shows the result of our generalization experiment.

### SUPPLEMENTARY VIDEOS

**Supplementary Video 1.** This video, which is available at https://youtube.com/shorts/RQ2torSVosM, shows a simulation run with chamois agents (white dots) and wolf agents (red dots) trained with the homeostatic reward signal.

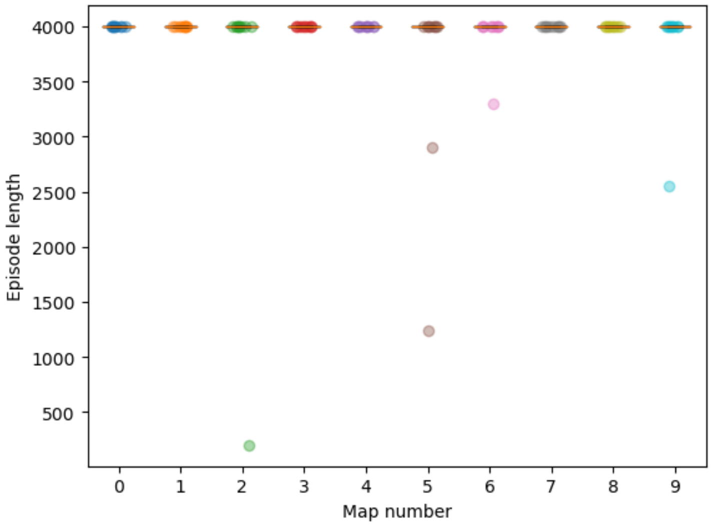


SUPPLEMENTARY FIGURE 3. Result of simulations on ten spatial models (Fig. 2). Ten simulations were run on each spatial model, starting from randomized agent distributions. The simulations (or episodes) lasted a maximum of 4,000 steps, but were aborted earlier if the wolves or chamois died out. Such extinction events happened in five of the 100 simulation runs.

**Supplementary Video 2.** This video, which is available at https://youtu.be/3rfpLHG2RjQ, shows a simulation run in a Perlin world with chamois and wolf agents controlled by a mix of policy networks. The policy networks either have two hidden layers and are trained with PPO, TOC, and TRPO, or no hidden layer and are trained with PPO, called PPO2.

**Supplementary Video 3.** This video, which is available at https://youtube.com/shorts/0SwnwIeSiBs, shows a simulation run of agents trained with the survival reward.

### SUPPLEMENTARY REFERENCES


[51] Annie Wong, Jacob de Nobel, Thomas Bäck, Aske Plaat, and Anna V Kononova. Solving deep reinforcement learning benchmarks with linear policy networks. *arXiv preprint arXiv:2402.06912*, 2024.

[52] Claes Strannegård, Niklas Engsner, Simon Ulfsbäcker, Sebastian Andreasson, John Endler, and Ann Nordgren. Survival games for humans and machines. *Cognitive Systems Research*, 86:101235, 2024.
